# Following the Trail of One Million Genomes: Footprints of SARS-CoV-2 Adaptation to Humans

**DOI:** 10.1101/2021.05.07.443114

**Authors:** Saymon Akther, Edgaras Bezrucenkovas, Li Li, Brian Sulkow, Lia Di, Desiree Pante, Che L. Martin, Benjamin J. Luft, Weigang Qiu

## Abstract

Severe acute respiratory syndrome coronavirus 2 (SARS-CoV-2) has accumulated genomic mutations at an approximately linear rate since it first infected human populations in late 2019. Controversies remain regarding the identity, proportion, and effects of adaptive mutations as SARS-CoV-2 evolves from a bat-to a human-adapted virus. The potential for vaccine-escape mutations poses additional challenges in pandemic control. Despite being of great interest to therapeutic and vaccine development, human-adaptive mutations in SARS-CoV-2 are masked by a genome-wide linkage disequilibrium under which neutral and even deleterious mutations can reach fixation by chance or through hitchhiking. Furthermore, genome-wide linkage equilibrium imposes clonal interference by which multiple adaptive mutations compete against one another. Informed by insights from microbial experimental evolution, we analyzed close to one million SARS-CoV-2 genomes sequenced during the first year of the COVID-19 pandemic and identified putative human-adaptive mutations according to the rates of synonymous and missense mutations, temporal linkage, and mutation recurrence. Furthermore, we developed a forward-evolution simulator with the realistic SARS-CoV-2 genome structure and base substitution probabilities able to predict viral genome diversity under neutral, background selection, and adaptive evolutionary models. We conclude that adaptive mutations have emerged early, rapidly, and constantly to dominate SARS-CoV-2 populations despite clonal interference and purifying selection. Our analysis underscores a need for genomic surveillance of mutation trajectories at the local level for early detection of adaptive and immune-escape variants. Putative human-adaptive mutations are over-represented in viral proteins interfering host immunity and binding host-cell receptors and thus may serve as priority targets for designing therapeutics and vaccines against human-adapted forms of SARS-CoV-2.

## Introduction

### Evolution in action: a trail of one million viral genomes

The 2002-2004 severe acute respiratory syndrome (SARS) coronavirus outbreaks had multiple origins (Chinese SARS Molecular Epidemiology Consortium 2004). In contrast, severe acute respiratory syndrome coronavirus 2 (SARS-CoV-2), the causative agent of the COVID-19 pandemic, showed nearly 100% sequence identity among the first outbreak strains from China, suggesting a single point of viral breach (Lu *et al*. 2020; Zhou *et al*. 2020). However, sequence diversity quickly accumulated as COVID-19 spread globally and remained uncontrolled a year later (Andersen *et al*. 2020; To *et al*. 2021). This high-stake case of evolution in action has brought unprecedented health, economic, and social devastation in modern times (Peeri *et al*. 2020; Kissler *et al*. 2020; To *et al*. 2021). Many of the evolutionary mechanisms driving SARS-CoV-2 genome diversification are unknown and urgently require elucidation. For example, to what extent has SARS-CoV-2 adapted to its new human hosts after one year of genome evolution (Phan 2020; Cagliani *et al*. 2020; Bai *et al*. 2020; Yang *et al*. 2020)? Moreover, how long can the global vaccine campaigns, most of which rely on vaccines formulated on basis of the bat-adapted viral genome, maintain effectiveness against the waves of new viral variants emerging worldwide (Koyama *et al*. 2020; Burton and Topol 2021)?

The vast number of publicly available SARS-CoV-2 genomes – expected to surpass a million before June 1, 2021 – offers unique opportunities for understanding the evolutionary processes accompanying the rapid emergence of a new viral pathogen, while challenging the ability to translate evolutionary understandings into the control and prevention of current and future pandemics (de Wit *et al*. 2016; Hadfield *et al*. 2018; Cui *et al*. 2019; Benvenuto *et al*. 2020; Andersen *et al*. 2020; Cagliani *et al*. 2020). Here we tested the hypothesis of rapid adaptation of SARS-CoV-2 genomes to human populations during the first year of the global COVID-19 pandemic. We focused on developing methods and computational tools for identifying human-adaptive mutations in the genomes of zoonotic viral pathogens. Identifying human-adaptive mutations is essential to uncovering the molecular mechanisms underlying the transition of SARS-CoV-2 from bat to human hosts, as well as the viral mechanisms of human pathogenesis and virulence (Cagliani *et al*. 2020). For disease treatment and prevention, human adaptive mutations are prime targets for the development of therapeutics against human-adapted SARS-CoV-2 variants as well as the development of broadly effective escape-proof vaccines (Burton and Topol 2021; Cohen *et al*. 2021).

### Challenges of identifying adaptive mutations in an asexual microbial population

Despite the benefits of a small genome size (~30,000 base pairs) and an abundance of geographically and longitudinally marked genome samples, identifying signatures of natural selection in SARS-CoV-2 is hindered by the challenge of a compact, gene-rich genome with few non-coding sequences, as is typical for microorganisms (DeLong 2004; Rocha 2018). Sequence evolution at non-coding loci in eukaryotic species hews closely to the standard Neutral Theory of molecular evolution, thus providing a powerful control for testing the presence of natural selection in functional genomic regions (Garud *et al*. 2015; Koropoulis *et al*. 2020). For example, presence of balancing (i.e., diversifying) selection at the *Adh* locus in *Drosophila* was discovered by an excess of nucleotide polymorphisms in the coding region relative to the 5’-flanking sequences (HKA test) (Hudson *et al*. 1987). The unexpected decrease in non-coding sequence diversity in genomic regions with low recombination rates has led to the discovery of pervasive purifying (i.e., negative) selection in *Drosophila* and humans (Hudson and Kaplan 1995; Charlesworth 2013; Campos and Charlesworth 2019). Likewise, adaptive mutations cause selective sweeps and reduce genetic diversity at linked non-coding loci (Sabeti *et al*. 2002; Garud *et al*. 2015).

Genome-wide linkage disequilibrium (LD) imposes an additional, more severe constraint for detecting adaptive mutations during SARS-CoV-2 evolution in human populations. In bacterial species, recombination is infrequent, yet it occurs at rates high enough to uncouple the evolution of loci under diversifying election (e.g., loci encoding surface antigens) from evolution of housekeeping loci under purifying selection (Milkman and Bridges 1990; Haven *et al*. 2011; Bobay *et al*. 2015). In sexual populations, proportions of adaptive amino-acid divergence could be estimated at a protein-coding locus by contrasting levels of synonymous and nonsynonymous substitution rates within and between species (MK test) (McDonald and Kreitman 1991; Charlesworth and Eyre-Walker 2006). However, the standard MK test severely underestimates adaptive divergence in asexual populations due to accumulation of slightly deleterious non-synonymous mutations (Charlesworth and Eyre-Walker 2008; Messer and Petrov 2013). Furthermore, both background selection and selective sweeps (“genetic draft”) reduce the effective population size and elevate the chance of random fixation of neutral and deleterious non-synonymous mutations in an asexual population, thus biasing the estimation of adaptive mutation rates (Gillespie 2000; Messer and Petrov 2013).

### Similarities between microbial experimental evolution and SARS-CoV-2 evolution

Experimental evolution under controlled laboratory conditions using microorganisms provides perhaps the most pertinent model for understanding SARS-CoV-2 evolution in humans (Lenski 2017; Good *et al*. 2017; Cvijović *et al*. 2018; Bergh *et al*. 2018). Although SARS-CoV-2 is a non-free-living organism evolving under open and diverse environmental conditions, SARS-CoV-2 populations share several key evolutionary characteristics with microbial populations in long term evolution experiments (LTEEs) (Lenski 2017; Cvijović *et al*. 2018). First, both evolving systems were seeded with a single genetically identical clone. Second, species in both systems were microorganisms containing a compact and gene-rich genome with few non-coding loci. Third, both systems had large populations in which natural selection was expected to prevail over genetic drift. For example, in a population with *N_e_* = 1000 individuals, any mutation with a selection coefficient |*s*| > 0.001 would cross the neutral barrier *N_e_s* = 1 and evolve deterministically towards fixation or extinction. Fourth, although capable of recombination, populations in both systems evolved clonally without detectable levels of genetic exchange among coexisting individuals. Thus, both systems evolved under genome-wide LD and were expected to show strong clonal interference (Lang *et al*. 2013; Lenski 2017; Good *et al*. 2017). Fifth, populations in both evolving systems were tracked in great genetic detail through whole-genome sequencing of temporally sampled isolates with spatial replication. Resembling the replicated populations in LTEEs, SARS-CoV-2 subpopulations in six continents (Asia, Africa, Europe, North America, South America, and Oceania) allowed for detection of adaptive changes based on recurring genetic events.

As expected given the strong similarities in key evolutionary characteristics, we found that SARS-CoV-2 populations during the COVID-19 showed similar adaptive dynamics as the *E. coli* populations in LTEE, including the early rise and rapid fixation of adaptive mutations, competing adaptive mutations, and recurrent genetic changes at key gene loci. Previous analyses of genome evolution of SARS coronaviruses have relied mainly on phylogenetic approaches to identify adaptive genes, haplotypes, and lineages (Chinese SARS Molecular Epidemiology Consortium 2004; Phan 2020; Cagliani *et al*. 2020; Bai *et al*. 2020; Yang *et al*. 2020). Crucially, without generating mutation spectra expected under neutral, background selection, and adaptive evolution models, these studies have been unable to test competing evolutionary models or to explore adaptive dynamics at the level of individual mutations. In LTEEs, the neutrally evolving populations were created by bottleneck events during serial transfer of cultures from one generation to another (Tenaillon *et al*. 2016). Here, we used *in silico* simulation of SARS-CoV-2 genomes evolving under neutral and selective models for understanding and predicting SARS-CoV-2 evolution during the COVID-19 pandemic.

## Material & Methods

### CoV genome simulator and the associated software system

*In silico* simulations are a powerful approach to test evolutionary hypotheses by providing fully specified evolutionary processes and parameters as models of species evolution in nature (Yuan *et al*. 2012). However, software tools for simulating the evolution of the gene-rich, finite-size microbial genomes such as those of SARS-CoV-2 are lacking. Simulations based on coalescent (backward-evolution) are highly efficient but are more suitable for modeling the evolution of neutral loci and relatively simple forms of selective and demographic mechanisms (Hudson 2002; Liang *et al*. 2007; Kelleher *et al*. 2016). Software tools based on forward-evolution simulations are less efficient but more flexible in modeling arbitrary selective and demographic forces (Carvajal-Rodríguez 2008; Hernandez 2008; Haller and Messer 2019). For simulating microbial genome evolution, two coalescent-based software tools implemented the realistic form of homologous recombination in bacterial genomes, but were not designed to simulate protein-coding sequences or the strong purifying and positive selective forces commonly operating on the gene-rich microbial genomes (Didelot *et al*. 2009; Brown *et al*. 2016). Furthermore, to our knowledge all existing simulation software implements infinite-site models of nucleotide substitutions. Consequently, these software tools do not allow for estimation of the chances of recurrent mutations at the same sites, an aspect that cannot be ignored in a rapidly expanding viral population with a small genome, such as SARS-CoV-2 populations during the COVID-19 pandemic.

Previously, we used forward-simulation to validate the origin and maintenance of high sequence diversity at a major surface antigen locus in the Lyme disease bacterium (*Borrelia burgdorferi*) by negative frequency-dependent selection (Haven *et al*. 2011). Here we developed a CoV genome evolution simulator (*CovSimulator*) and used it to test whether patterns of CoV genome variability fit better with expectations from neutral (NEU), background-selection (BKG), adaptive (ADPT), or mixed (MIX) evolution models. The software system associated with the *CovSimulator* is diagramed in Supplemental Material Fig S1.

Briefly, *CovSimulator* first read the annotated genome of a viral progenitor provided in GenBank format (e.g., Wuhan-Hu-1, GenBank accession NC_045512). It captured the reading frames of the 25 protein-coding loci (Table 1) in the SARS-CoV-2 genome such that coding (or non-coding) information associated with each base of the genome was stored. At a protein-coding nucleotide site, the stored genomic information included the gene locus, codon, amino acid, and codon position. Simulation was initialized with a population of *N* identical ancestral genomes, each of which was assigned the unit fitness value (see evolution parameters in Table 2). During each of the total number of *g* generations, each individual encountered a Poisson-distributed number of point mutations with the mean genome mutation rate *m*. If a mutation occurred, a uniformly distributed genome position was chosen and an alternative nucleotide was selected as the substitute according to the base-substitution frequencies gathered from viral genomes (see section below). Similarly, homologous recombination during each generation occurred with a Poisson distributed mean rate of *r* per genome. If a recombination event occurred, two individuals from the population were randomly chosen and a uniformly distributed genome position was selected as the break point. Two new individual genomes were created by exchanging the sequences right and left of the break point. Fitness values of the new genomes were re-computed according to a new set of mutated sites.

**Table 1.** Protein-coding loci (*n*=25) included in simulated SARS-CoV-2 genome evolution

| Protein ID<sup>a</sup> | Locus symbol | Protein function<sup>b</sup> | Location<sup>a</sup> | Locus length | Syn sites (S)<sup>c</sup> | Missense sites (N)<sup>c</sup> |
| --- | --- | --- | --- | --- | --- | --- |
| YP_009725297.1 | *nsp1* | Leader protein; inhibits host translation | 266, 805 | 540 | 164.48 | 375.52 |
| YP_009725298.1 | *nsp2* | Unknown | 806, 2719 | 1914 | 583.94 | 1330.06 |
| YP_009725299.1 | *nsp3* | Polyprotein processing | 2720, 8554 | 5835 | 1738.36 | 4096.64 |
| YP_009725300.1 | *nsp4* | Formation of double membrane vesicles associated with replication complexes | 8555, 10054 | 1500 | 449.50 | 1050.50 |
| YP_009725301.1 | *nsp5* | 3C-like proteinase; polyprotein processing | 10055, 10972 | 918 | 278.19 | 639.81 |
| YP_009725302.1 | *nsp6* | Formation of double membrane vesicles associated with replication complexes | 10973, 11842 | 870 | 253.34 | 616.66 |
| YP_009725303.1 | *nsp7* | Accessory subunit of RNA-dependent RNA polymerase | 11843, 12091 | 249 | 75.97 | 173.03 |
| YP_009725304.1 | *nsp8* | Accessory subunit of RNA-dependent RNA polymerase; primase | 12092, 12685 | 594 | 164.77 | 429.23 |
| YP_009725305.1 | *nsp9* | RNA-binding protein | 12686, 13024 | 339 | 105.61 | 233.39 |
| YP_009725306.1 | *nsp10* | Co-factor of Nsp14 and Nsp16 for methyltransferase activity | 13025, 13441 | 417 | 123.76 | 293.24 |
| YP_009725307.1 | *nsp12* | RNA-dependent RNA polymerase | 13442, 13468; 13468, 16236 | 2796 | 843.60 | 1952.40 |
| YP_009725308.1 | *nsp13* | Helicase | 16237, 18039 | 1803 | 541.37 | 1261.63 |
| YP_009725309.1 | *nsp14* | Proof-reading 3'-to-5' exonuclease | 18040, 19620 | 1581 | 469.80 | 1111.20 |
| YP_009725310.1 | *nsp15* | Endonuclease | 19621, 20658 | 1038 | 319.98 | 718.02 |
| YP_009725311.1 | *nsp16* | Ribose 2'-O-methyltransferase; RNA cap formation | 20659, 21552 | 894 | 268.86 | 625.14 |
| YP_009724390.1 | *S* | Surface glycoprotein; spike protein; binding of host cell receptor | 21563, 25384 | 3822 | 1150.89 | 2671.11 |
|  | *S1* | S1 subunit containing the angiotensin- converting enzyme 2 (ACE2) receptor-binding domain (RBD) (Wang *et al.* 2020; Huang *et al.* 2020) | 21563, 23185 | 1623 | 482.32 | 1140.68 |
|  | S2 | S2 subunit responsible for membrane fusion | 23186, 25381 | 2296 | 667.38 | 1528.62 |
| YP_009724391.1 | *orf3a* | Ion channel promoting virus release (Lu *et al.* 2006; Siu *et al.* 2019) | 25393, 26220 | 828 | 252.54 | 575.46 |
| YP_009724392.1 | *E* | Envelope protein forming homopentameric cation channel | 26245, 26472 | 228 | 80.43 | 147.57 |
| YP_009724393.1 | *M* | Membrane glycoprotein | 26523, 27191 | 669 | 202.40 | 466.60 |
| YP_009724394.1 | *orf6* | Unknown | 27202, 27387 | 186 | 53.74 | 132.26 |
| YP_009724395.1 | *orf7a* | Unknown | 27394, 27759 | 366 | 115.71 | 250.29 |
| YP_009725318.1 | *orf7b* | Unknown | 27756, 27887 | 132 | 46.24 | 85.76 |
| YP_009724396.1 | *orf8* | Ion channel contributing to suppression of host immunity (Zinzula 2021) | 27894, 28259 | 366 | 121.59 | 244.41 |
| YP_009724397.2 | *N* | Nucleocapsid phosphoprotein; viral RNA genome protection and packaging | 28274, 29533 | 1260 | 396.51 | 863.49 |
| YP_009725255.1 | *orf10* | Unknown; suspected membrane protein forming viroporin | 29558, 29674 | 117 | 32.88 | 84.12 |

<sup>a</sup> Start and stop positions (stop codon included) based on the genome of the strain Wuhan-Hu-1 (GenBank Accession NC\_045512) (Zhou *et al.* 2020). The *nsp11* locus (39 bases; 13442-13480), which is small and enclosed within the *nsp12* locus, was excluded. The 4-base overlap between *orf7a* and *orf7b* was counted towards the former locus.
<sup>b</sup> (To *et al.* 2021)
<sup>c</sup> Effective numbers of synonymous (*S*) and nonsynonymous (*N*) substitution sites of a protein-coding locus, derived with the *CovSim-* *ulator* (see Material and Methods)

Crucially, we defined the fitness of a simulated viral genome as its adaptiveness to the human host relative to the ancestral viral genome. That is, the fitness of the ancestral viral genome to the human host was defined as one. A simulated viral population displaying an average fitness > 1 could thus be interpreted as being better adapted than the ancestral genome to the human host. A simulated viral population with an average fitness = 1 was considered equally fit as the ancestral viral genome to reproduce in the human host. Otherwise, a simulated viral population with an average fitness < 1 was considered less fit than the ancestral genome to use the human host. To implement this fitness scheme, we determined synonymous or missense mutations and computed the fitness value of a simulated genome according to comparison with the ancestral viral genome rather than its parental genome. This fitness definition is equivalent to measuring fitness gains in an LTEE study through competing the evolved strains with the original, pre-evolved strain (Lenski 2017).

The fitness of an individual genome was the multiplicative product of its composite codons. Thus, the fitness of the individual was set to zero if the mutation introduced a stop codon (nonsense) or changed a stop codon into a sense codon (reading-frame extension). Otherwise, the mutation introduced an amino-acid change (missense mutation). In the neutral model, the fitness of an individual remained unchanged by missense mutations. In the background selection model, a missense mutation had a probability of *u* (e.g., *u* = 0.8) of decreasing the fitness of its carrier genome by a factor of, e.g., *w* = 0.95. In the adaptive evolution model, in contrast, a missense mutation had a small probability of *v* (e.g., *v* = 0.1) of increasing the fitness of its carrier genome by a factor of, e.g., *w* = 1.05. The fitness of the individual was unchanged if the mutation occurred at a non-coding (intergenic or untranslated) site or introduced a synonymous amino acid.

The probability of an individual to produce an offspring in the next generation was determined by its fitness. Specifically, a threshold value between 0 and 1 was computed as an increasing function of the fitness of an individual *c* = 1 – *e*^−*w*^. A random number *p* between 0 and 1 was chosen. If *p* < *c*, then the individual was able to contribute one offspring. Otherwise, the individual did not have a chance to reproduce. The parental population was repeatedly sampled with replacement for reproduction until the constant population size of *N* was reached. To validate the genome simulator, we compared the sample statistics with neutral expectations including the level of sequence polymorphism at mutation-drift balance (*θ* = 2*N_e_μ*_0_), the rate of sequence divergence with respect to the ancestor (*k*=*mt*), and the length and shape of genome genealogies under neutral and selective evolution.

### Viral genome database and the associated software system

Viral genomes and associated information on the geographic origins and collection dates were obtained from GISAID monthly according to submission dates (Shu and McCauley 2017). SNVs and indels in each genome with respect to the reference genome (Wuhan-Hu-1, GenBank accession NC_045512) were identified by using the program DNADIST in the Nu-cmer4 package (Marçais *et al*. 2018). To minimize sequencing errors, SNVs at genome ends where missing bases were common were excluded, as were any genomes with more than 10% missing bases at SNV sites. Unique haplotypes were obtained with custom Perl scripts based on the BioPerl package (Stajich *et al*. 2002). Isolate information, variants, and haplotypes were deposited into a custom relational database (“cov-db”) to facilitate downstream computational analysis. A custom Python script sampled viral genomes (e.g., *n*=100) by month and at three spatial scales (continent, country, and state). The script also filtered variants and output only the most frequently occurring (e.g., >0.5%) variants. A secondary Python script produced a variant call format (VCF) file based on the sampled isolates and high-frequency variants. Evolutionary statistics, including variant frequencies, linkage disequilibrium (*r^2^*), haplotypes, and base substitution frequencies were generated with programs BCFTools and VCFTools (Danecek *et al*. 2011). We used Haploview (version 4.2) to calculate LD scores (*D*’ and *r^2^*) as well as their statistical significance between pairs of SNVs (Barrett 2009). The most parsimonious haplotype networks were estimated with the program TCS ver1.21 (Clement *et al*. 2000) and visualized with tcsBU (Múrias dos Santos *et al*. 2016). To visualize genome variants and haplotype networks, we developed a custom web interface (http://genometracker.org) using a similar software system supporting *BorreliaBase*, a comparative genomics browser of Lyme disease pathogens (Di *et al*. 2014). The software system associated with the “cov-db” database is diagramed in Supplemental Material Fig S2.

### Evolution rates, linkage disequilibrium, and homoplasy

We estimated the SARS-CoV-2 genome divergence rate from the ancestor by performing a linear regression of sequence differences to the reference genome (NC_045512) with respect to the genome collection dates. The expected variance of the evolutionary rate was estimated according to a Poisson model, which specified, at each time point of *t* days, an expected number of sequence differences in *λ*=*μtL*, where *μ* being the rate of base substitution per site per day obtained from the regression line and *L* being the length of the reference genome (NC_045512, *L*=29903). The variance of the Poisson expected difference was expected to be equal to the difference itself (*σ^2^=λ*).

To compare cross-species rates of amino-acid substitutions at protein-coding loci, we downloaded the genomes of 24 viral isolates belonging to the family Coronaviridae. The viral isolates included coronaviruses closely related to SARS-CoV and SARS-CoV-2 and consisted of Wuhan-Hu-1, RaTG13, P1E, P5L, ZC45, ZXC21, SC2018, HuB2013, Shaanxi2011, HKU3-1, Rm1, CoV273, GX2013, Rf4092, YN2013, GD01, SZ3, WIV16, SHC014, YN2018B, As6526, Rs4247, Rs672, and Yunnan2011. Homologous protein sequences were aligned and individual alignments were concatenated using the sequence utility *bioaln* from the BpWrapper software suite (Hernández *et al*. 2018). Per-site substitution rates, normalized to a mean rate of zero, were obtained with *rate4site* (Pupko *et al*. 2002).

We used Haploview (version 4.2) to calculate linkage disequilibrium (LD) scores (*D*’ and *r^2^*) as well as their statistical significance (LOD, log odds) between pairs of SNVs (Barrett 2009). We used the DNAPARS program of the PHYLIP (version 3.696) package to search for a maximum parsimony tree of unique haplotypes, obtaining the homoplasy index (HI) and the number of base substitutions at each SNV site (Felsenstein 1989). The HI is defined as 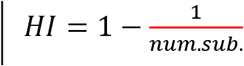 at each SNV site and is zero when the alleles are consistent with the tree (i.e., the number of substitution for a bi-allelic SNV is one).

### Analysis of synonymous and missense evolutionary rates

Genome-wide numbers of synonymous (*D_s_*) and nonsynonymous (*D_n_*) nucleotide divergence were obtained through comparison of the reference genome (Wuhan-Hu-1, GenBank accession NC_045512) to its closest known relative (RaTG13, GenBank accession MN996532) (Zhou *et al*. 2020) with the program DNADIST (Marçais *et al*. 2018). Genome-wide synonymous (*P_s_*) and nonsynonymous (*P_n_*) nucleotide polymorphisms in viral populations were estimated with the use of viral samples. In computing the per site synonymous and nonsynonymous substitution rates (*d_s_*=*D_s_/S, d_n_=D_n_/N, p_s_=P_s_/S, p_n_=P_n_/N*), the effective numbers of available synonymous (*S*) and nonsynonymous (*N*) sites at each gene locus must be estimated (Yang 2007). We estimated *S* and *N* empirically by using the *CovSimulator*, which accounts for both the genome base composition bias and the strong mutation biases (Supplemental Material Fig S3). Specifically, we ran CovSimulator with a high genome mutation rate *m*=10 and a population size *p*=200 for *n*=10 generations, generating an expected total number of 20,000 mutation events or *λ*=0.67 mutations per genomic site, on average. Assuming a Poisson distribution, the proportions of genomic sites encountering 0, 1, and >1 point mutations were expected to be 51.2%, 34.3%, and 14.5%, respectively. Thus, the probability of a site not being mutated was *p*=0.512. To ensure that all genomic sites were mutated at least once, we ran *CovSimulator* ten times such that the chance of a site not undergoing any mutation was small *p* = 0.512^10^ = 1.25e-3. The average numbers of synonymous and missense mutations from ten repeated runs, normalized to gene lengths, were used as estimates of *S* and *N* (Table 1; Supplemental Material Tables S1 and S2).

### Analysis of mutation trajectories

For a simulated population, we followed the trajectories of the most frequent (>0.5% among all samples) missense mutations by first calculating their frequencies in each generation. The trajectory of a mutation *X* was represented by an *n*-dimensional vector *X_T_* = (*X*_*t*_0__,…., *X_t_n__*), where each *x_t_i__* was the frequency of *X* within in the population at time point *t_i_*. Distance between two trajectories, *X_T_* and *Y_T_*, was defined as 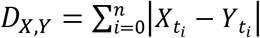. Trajectories of two or more mutations were merged into a “genotype” if the average distance between them was ≤ 0.05. A genotype (*G_1_*) was considered as derived from (i.e., nested within) a parental genotype (*G_2_*) if their Jaccard distance 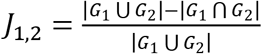 equaled the simplified Jaccard distance when *G_1_* was nested within *G_2_*, 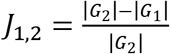. In the latter case, the union of the two trajectories |*G*_1_ ∪ *G*_2_| was the same as the trajectory of the parental genotype *G*_2_, whereas the intersection of the two trajectories |*G*_1_ ∩ *G*_2_| was the same as the trajectory of the child genotype *G*_1_. The merging of mutations into genotypes and the nesting of genotypes were both carried out with the Python package *muller* (version 0.6.0, https://github.com/cdeitrick/Lolipop) with default settings. Muller diagrams were subsequently generated using the R package *ggmuller*. Fitness of a “genotype” was defined cumulatively (Desai and Fisher 2007). An adaptive mutation within a cluster increases its fitness by *s*>0 and a deleterious mutation decreases its fitness by *s*<0. If a parent cluster had fitness *ns*, and a child of that cluster had fitness *ms*, the fitness of the child was (*n+m*)*s*.

For viral genome samples, Muller diagrams, which depict mutation frequencies within a single evolving population, are in general not applicable. Since viral genomes were sampled from multiple outbreak locations, it would be misleading to perform analysis including merging of mutations into genotypes and inference of parental and child genotypes. Thus, we used heatmaps as an alternative approach to follow the trajectories of high-frequency mutations in both simulated and viral populations. The R package *pheatmap* was used to generate heatmaps. As in the Muller diagrams, mutations in a heatmap were grouped into hierarchical clusters based on similarities in frequencies over time. Similarity between a pair of mutation trajectories *i*, and *j* was defined as *d* = 1 – *cor*(*i, j*), where *cor*(*i, j*) was the Pearson’s correlation coefficient. Unlike in the Muller diagrams, however, mutations with similar frequency trajectories were not merged into “genotypes”. Nor did the heatmap analysis estimate parent-descendant relationships among mutation clusters.

### Data and software availability

SARS-CoV-2 genome sequences and the associated viral isolate information are available from the GISAID EpiCoV™ database (Shu and McCauley 2017). Software tools associated with *CovSimulator*, the forward-evolution simulator, and *cov-db*, the custom database of SARS-CoV-2 genome variability, are available at the Github repository (https://github.com/weigangq/cov-db). Programmatic access to the *cov-db* database is available upon request. Also available in the same Github repository are key datasets including mutation trajectories from simulated evolution and VCF files of viral genomes sampled monthly. A web interface to the *cov-db* database is publicly available at http://cov.genometracker.org.

## Results and Discussion

### *CovSimulator*. a realistic SARS-CoV-2 genome evolution simulator

The SARS-CoV-2 genome is biased in base composition (62.0% AT) and strongly biased in mutation frequency. Approximately ~70% of single-nucleotide mutations occurring during the pandemic were C>T or G>T substitutions (Supplemental Material Fig S3). To realistically simulate SARS-CoV-2 genome evolution, we used the first known SARS-CoV-2 genome (from the Chinese isolate Wuhan-Hu-1 collected in December 2019) (Zhou *et al*. 2020) as the progenitor and used empirically derived base substitution probabilities (Table 2). *CovSimulator*, currently implemented with the standard Wright-Fisher model with constant population sizes and non-overlapping generations, was validated through comparing simulated outputs with analytical expectations including the rates of sequence divergence (Fig 1C and 1D), levels of sequence polymorphism (Fig 2C), genealogies of genome samples at the end of simulations (Supplemental Material Fig S4A), and fitness values of simulated populations (Supplemental Material Fig S4B).

**Fig 1.**
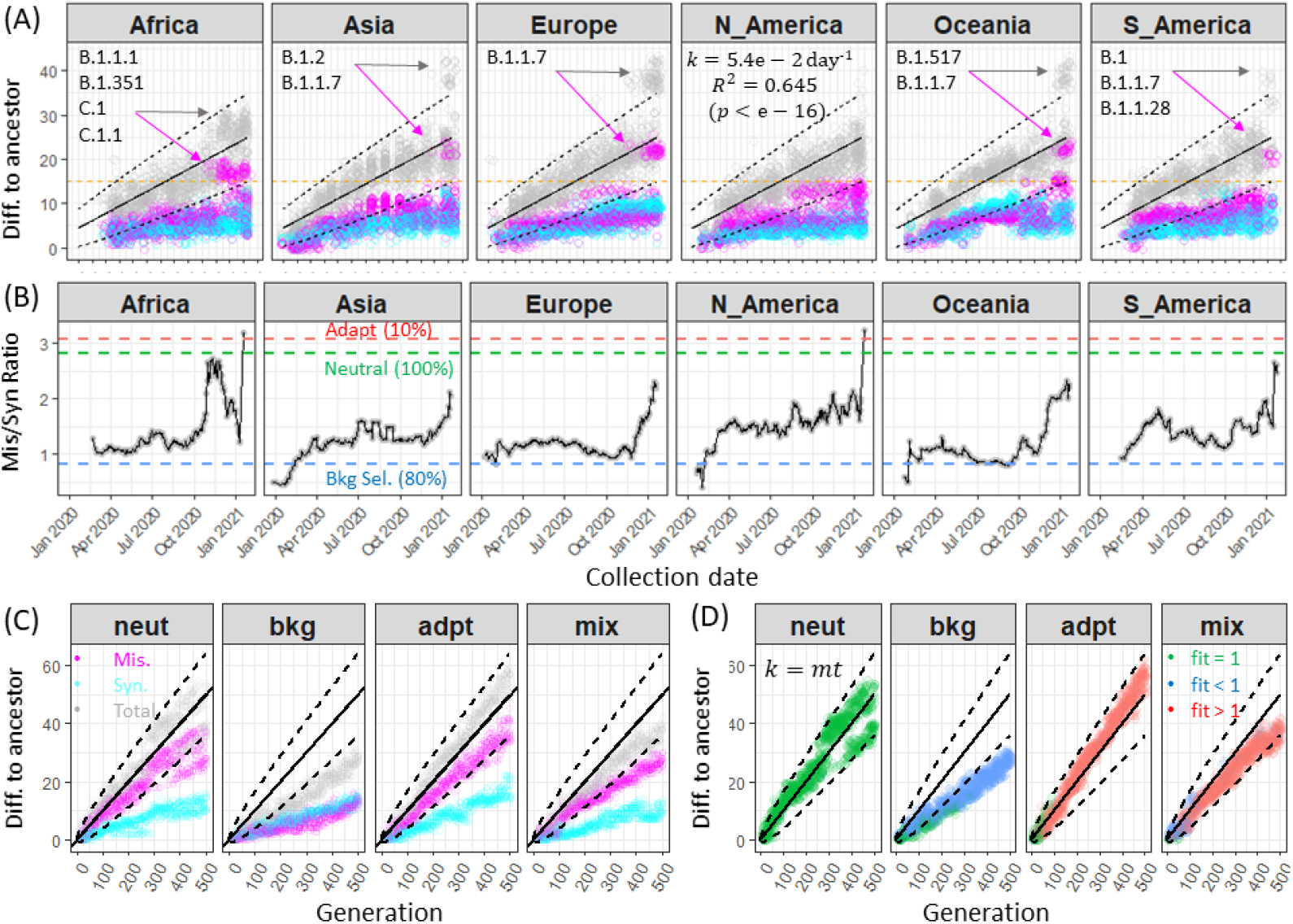
Rates of synonymous and nonsynonymous divergence of SARS-CoV-2 genomes. (A) In each panel, points represent differences with respect to the reference genome (*y*-axis) of viral genomes originating in a continent with collection dates (*x*-axis) ranging from Dec 2019 through March 2021. A random sample of 100 genomes was chosen for each month, resulting in ~1400 genomes for each continent with evenly distributed monthly representations. Each genome was represented three times, including the number of synonymous mutations (cyan), the number of missense mutations (magenta), and the total number of genetic changes (including indels; gray). A linear regression line (solid, with statistics shown within the “N_America” panel) was derived by using genomes from all continents. Dashed lines show two standard deviations above and below the regression line on the basis of the Poisson expected variance *σ*^2^(*k*) = *k*. Hyper-mutated genomes (marked by the lineage designations) that emerged in late 2020 showed accelerated accumulation of missense (but not synonymous) mutations (Choi *et al*. 2020; Kemp *et al*. 2021). The orange lines indicate a cutoff value of 15 missense mutations to determine outliers. (B)Ratios of the numbers of missense (*D_n_*) to synonymous (*D_s_*) mutations relative to the reference genome (*y*-axis), a measure of selective constraint, were plotted against the viral sample collection dates (*x*-axis). Each point was the ratio of the average number of missense to synonymous mutations within a moving window of 14 days. Horizontal dashed lines mark the ratios obtained from simulated evolution and percentages represent proportions of missense mutations that were deleterious (blue), neutral (green) and adaptive (red) (see below). *D_n_/D_s_* ratios in all populations started at low levels, indicating strong purifying selection during the early months (before April 2020) of the pandemic. The *D_n_/D_s_* ratio increased greatly in later months of 2020 worldwide, suggesting rapid population expansion and the emergence of human-adaptive viral variants. (C) Mutational divergence (*y*-axis) over the generation (*x*-axis) of genomes simulated with four evolution models. For each model, a sample of *s*=20 genomes was chosen for each generation, resulting in ~10,000 genomes for each model. Solid lines indicate the expected mutation rate in the neutral model. Dashed lines show two standard deviations above and below the expected total mutation rate on the basis of the Poisson expected variance *σ*^2^(*k*) = *mt*. Genome evolution was simulated with a population size *N=*200, genome mutation rate *m*=0.1 and no recombination (*r*=0). In the neutral evolution model (“NEUT”), all missense mutations carried a fitness of 1. In the background selection model (“BKG”), a missense mutation had an 80% chance of incurring a fitness cost of 0.95 and was otherwise neutral. In the adaptive evolution model (“ADPT”), a missense mutation had a 10% chance of incurring a fitness benefit of 1.05 and was otherwise neutral. In the mixed evolution model (“MIX”), a missense mutation had an 80% chance of incurring a fitness cost of 0.95, a 10% chance of incurring a fit benefit of 1.05, and 10% chance of being neutral. As expected, the ratio of missense to synonymous mutations (~1.0) in the BKG model was markedly lower than that in neutral evolution and was used as a control (blue dashed line in *panel B*) for measuring viral evolution during the pandemic. The ratios of missense to synonymous mutations from the neutral and mixed evolution models were much higher (~3.0) and were used as another set of controls (green and red dashed lines in *panel B*) to understand SARS-CoV genome evolution. (D) Simulated genomes colored by fitness values. In the neutral evolution model, the total mutation rate and its variability accurately followed the expectations, thereby validating the *CovSimulator*. In the background selection model, the overall mutation rate decreased and the population was increasingly dominated by low-fitness genomes, showing a gradual loss of fitness in a clonal population known as Muller’s ratchet (Muller 1964). In the adaptive evolution model, mutation accumulation was accelerated and the population was dominated by adaptive lineages except in the first 100 generations. In the mixed evolution model, adaptive lineages dominated the population despite the presence of strong purifying selection.

**Table 2.**
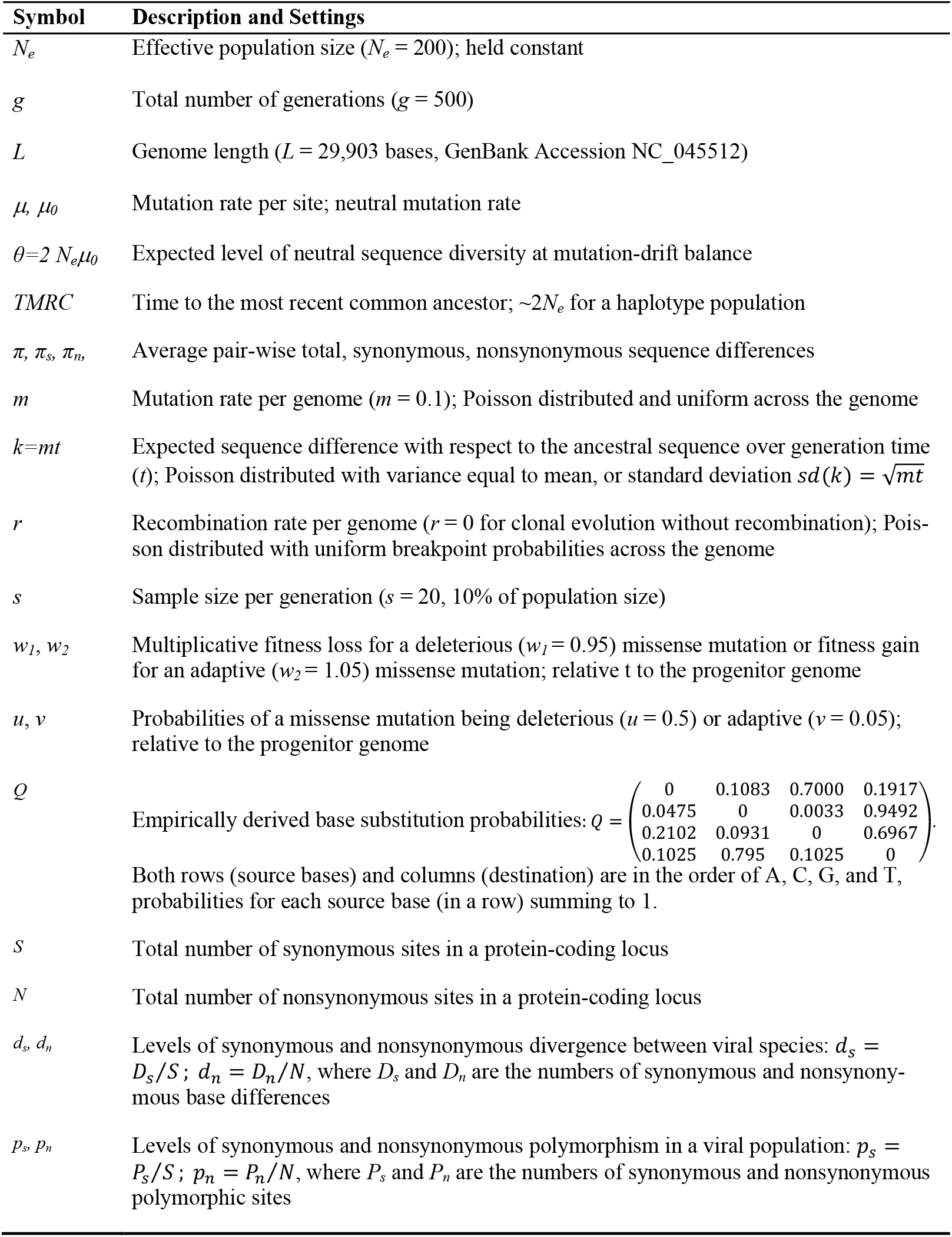
Evolutionary parameters and neutral expectations

We used the CovSimulator to derive theoretical expectations of synonymous and missense divergence, sequence diversity, and their ratios under neutral, background, adaptive, and mixed evolution models (Figs 1C, 1D, 2C, and 2D). These simulated outcomes provided baseline controls for estimating selective constraints (Figs 1B & 2B) as well as for understanding evolutionary dynamics at the level of individual mutations (Fig 3). In addition, the CovSimulator provided a simulation-based approach to estimate evolutionary parameters such as the effective numbers of synonymous and nonsynonymous sites at protein-coding loci (Table 1) and frequencies of recurrent mutations (see below). Such parameters are difficult to derive analytically because it is necessary to take into account of biases in base composition as well as in mutation frequency.

**Fig 2.**
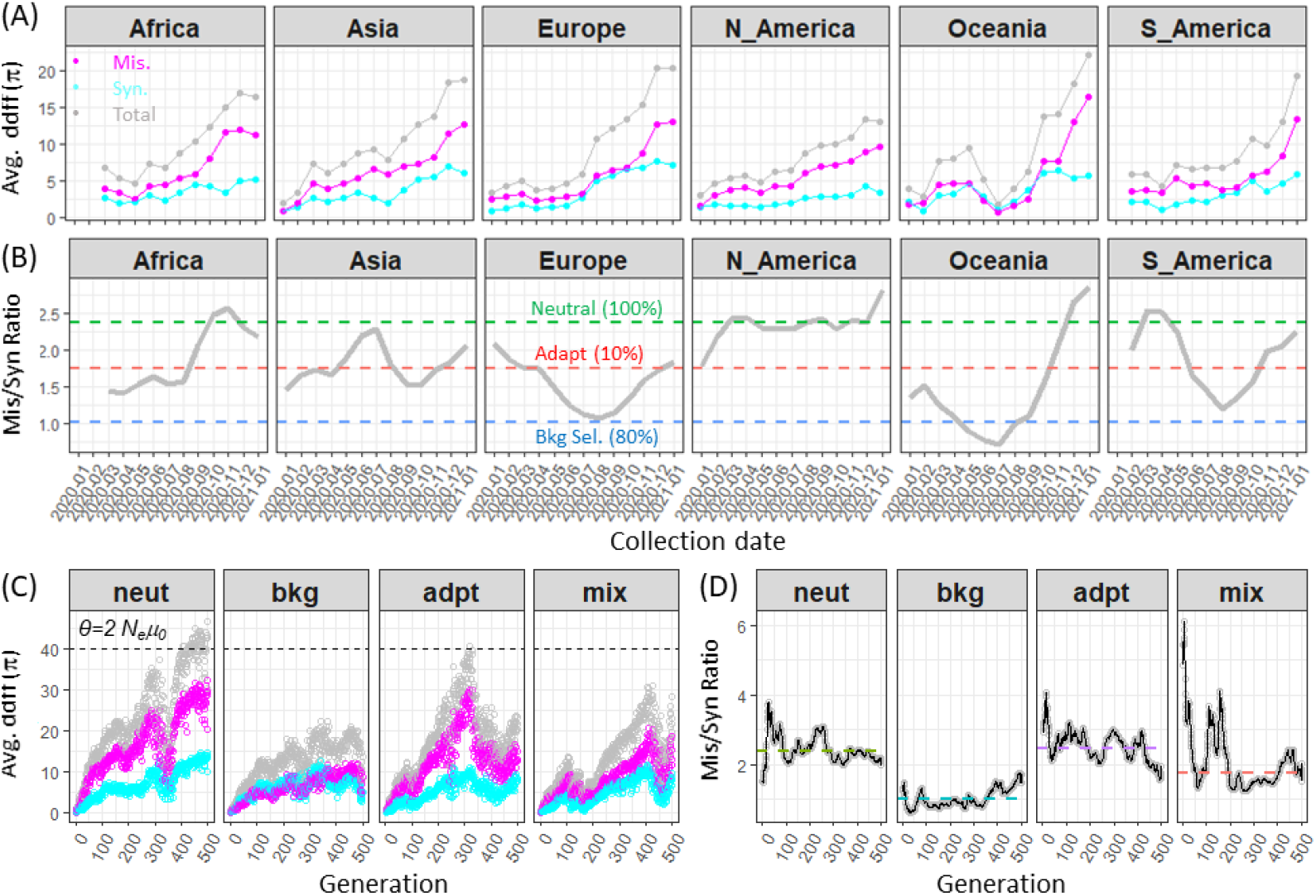
Synonymous and nonsynonymous polymorphisms of SARS-CoV-2 populations. The average number of pairwise sequence differences (*π*) is a measure of viral genetic diversity, which reflects the viral effective population size and viral reproduction rate. (A) Each panel represents a continental population. Synonymous (*π_s_*), nonsynonymous (*π_n_*), and total pairwise sequence differences (*y*-axis) were calculated from monthly samples from December 2020 through March 2021 (*x*-axis). Sequence diversity increased in all populations. An initial increase in genetic diversity was expected for a nascent viral population before reaching mutation-drift balance. However, the acceleration of viral diversity in later months was unexpected and reflected the accumulation of neutral, deleterious, and adaptive genetic diversity in rapidly expanding viral populations. (B) The ratio of nonsynonymous (*π_n_*) to synonymous (*π_s_*) diversity, similarly to *D_n_/D_s_* (Fig 1), is a measure of selective constraints. The *π_n_/π_s_* ratios were elevated and fluctuated substantially, in agreement with a lack of selective constraints in rapidly expanding viral populations. (C) The *π* values of simulated genomes under four evolution models. In the neutral evolution model the total *π* value stabilized at the expected value of *θ* = 40, further validating the *CovSimulator*. The *π* values were relatively lower in the background and adaptive selection models, in agreement with smaller effective population sizes due to natural selection and shorter coalescent times (Supplemental Fig S4). Genome evolution was simulated with a population size *N*=200, genome mutation rate *m*=0.1 and without recombination (*r*=0). In the neutral evolution model (“NEUT”), all missense mutations carried a fitness of one. In the background selection model (“BKG”), a missense mutation had an 80% chance of carrying a fitness cost of 0.95. In the adaptive evolution model (“ADPT”), a missense mutation had a 10% chance of carrying a fitness benefit of 1.05. (D) The *π_n_/π_s_* ratios for the four evolution models, showing a low value for a population under purifying selection, a high value during neutral evolution, and intermediate values for a population under adaptive evolution. These average ratios are shown in panel (B) as references for comparison with values based on viral samples.

**Fig 3.**
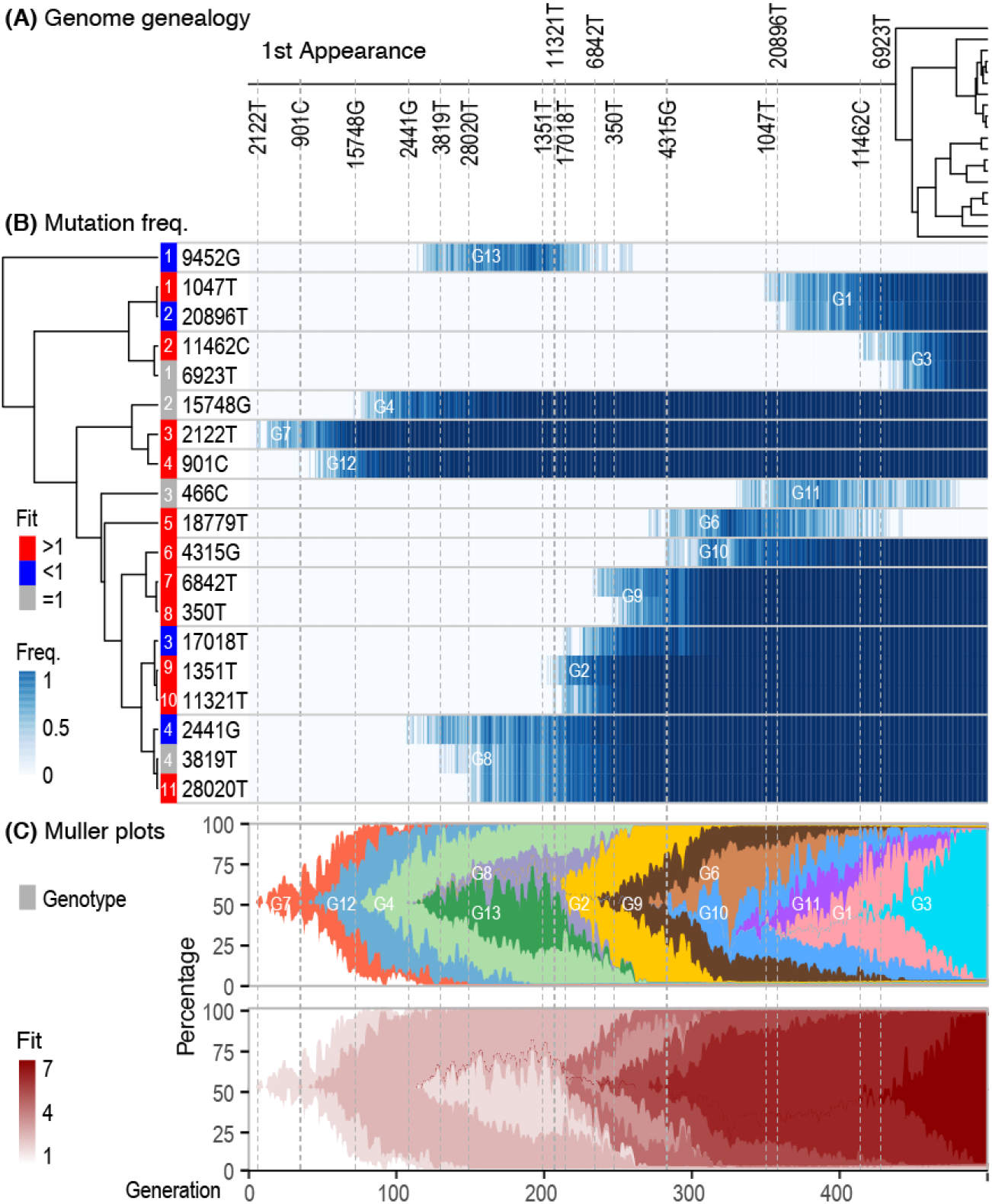
Mixed evolution as a model of SARS-CoV-2 genome evolution. Simulated evolution of a viral population subject to both purifying and adaptive selection (parameters defined in Fig 1) captured viral adaptation at the level of individual mutations. (A) Genealogy of the *n*=20 simulated genomes sampled from the last generation. The 16 mutations were fixed in the final population and their times of first appearance in the population (tick marks on the root edge) corresponded to the timing of selective sweeps shown in the Muller diagram (*panel C*). The genealogy shows that the final viral population descended from the latest selective sweep, which occurred only ~50 generations ago. (B) In the heatmap, rows show *n*=19 high frequency (≥5% in 10,000 genome samples) non-synonymous SNVs, labeled by the genome position and destination base, from the mixed evolution model. Frequencies were represented by color intensity (*middle panel*). Unlike those of natural viral variants, the fitness values of simulated variants were precisely known (*colored side bar*). The variants were grouped according to the correlated evolutionary trajectories (*dendrogram*) such that mutations with similar trajectories (“genotypes”) – indicating temporal linkage – were adjacent. Adaptive mutations (red, numbered from #1 through #11) dominated the population. Adaptive mutations (#3 and #4) arose early. One adaptive mutation (#5) was lost, probably because of clonal interference but also possibly because of genetic drift. Among the four deleterious mutations, one (#1) was lost whereas the other three hitchhiked with linked adaptive mutations to fixation. Similarly, one neutral mutation (#3) was lost and three others hitchhiked to fixation. Furthermore, the simulation suggests a high rate of multiple mutations occurring at the same genomic site associated with adaptive mutations (#1, #10, and #11). Three of the four multiple-hit mutations had T as the destination base, reflecting the strong mutation bias in which ~70% SNVs during viral evolution were due to C>T or G>T substitutions. (C) Muller diagrams of mutation trajectories in the simulated population. In the top diagram, at each vertical cross section, the heights of colored blocks represent the relative frequencies of the “genotypes”. A genotype represented one or more mutations displaying a similar evolutionary trajectory reflecting temporal linkage. The Muller diagram reveals selective sweeps occurring regularly and strong competition among adaptive mutations. For example, during generations 100 to 250, the top Muller plot shows competition between two genotypes (G8 and G13), both of which carried a deleterious mutation. They had similar fitness values (bottom Muller plot). At the ~250 generation, however, G13 was outcompeted by G8 and went extinct. G8 itself subsequently gave rise to G2. Similarly, at the ~300^th^ generation, two adaptive mutations (#5 in G6 and #6 in G10) began to compete against each other and coexisted until the ~450^th^ generation when G6 was eventually displaced by G1 and G3, both descendants of G10. The latter case represents a loss of an adaptive allele (#5) due to clonal interference.

Future versions of the simulator will incorporate more realistic demographic features including changing population sizes, population admixture, and additional selective mechanisms. In particular, simulating SARS-CoV-2 genome evolution under negative frequency-dependent selection is important to identify mutations contributing to immune escape including escape from vaccines, which are expected to have higher fitness values when they rare (Haven *et al*. 2011; Papkou *et al*. 2019). Negative frequency-dependent mutations maintaining coexistence of multiple clonal lineages have been observed in microbial LTEEs (Maddamsetti *et al*. 2015; Good *et al*. 2017). In addition, the CovSimulator paves a way to estimate parameters of SARS-CoV-2 evolution (e.g., population growth rates, selection coefficients, and migration rates) through approximate Bayesian computation (ABC) (Lintusaari *et al*. 2017).

### Accelerated missense divergence: rise of hyper-mutated variants

On the basis of ~1.0 million SARS-CoV-2 genome sequences obtained from the GISAID database (Shu and McCauley 2017) up to March 31, 2021, we generated a custom database of high-quality 815,402 SARS-CoV-2 genomes. We identified 8065 SNVs, 173 deletions, and 49 insertions, each of which was represented by 100 or more viral genomes. Genome sequences were consolidated into 350,094 haplotypes based on the SNVs and indels. We sampled ~100 genomes monthly from each of the six continental populations and plotted the synonymous, missense, and total mutational differences with respect to the ancestral genome over collection dates (Fig 1A). The total rate of mutation accumulation (gray dots) was well characterized by a linear Poisson model with a highly significant slope and a Poisson-expected variance (Fig 1A). However, significant deviations from the Poisson model were observed in Asian, European, Oceanian, and South American viral populations since October 2020, associated with the hyper-mutated viral variants discovered first in immuno-compromised patients with COVID-19 (Choi *et al*. 2020; Kemp *et al*. 2021).

The Poisson model of linear mutation accumulations over time initially suggested a neutral process of viral genome divergence. However, closer examination by measuring the synonymous and missense mutation rates separately did not support the neutral divergence model. The ratio of missense to synonymous mutations was expected to be high (*D_n_/D_s_*~3.0) according to the simulated neutral evolution (Fig 1C, *1^st^* panel). In reality, the *D_n_/D_s_* ratios began at a low level (*D_n_/D_s_*~1.0) similar to that from the simulated background selection model (Fig 1C, *1^st^* panel), thus suggesting considerable selective constraints during the early months (before April 2020) of the pandemic. The *D_n_/D_s_* ratio increased across continental populations afterward and eventually showed a marked increase after October 2020 to the levels expected from the neutral and adaptive evolution models (Fig 1B). The accelerated missense divergence, reflected in the steep rise of *D_n_/D_s_* ratios, was attributable to the emergence and spread of hyper-mutated viral lineages, which accumulated predominantly missense mutations with little synonymous divergence (Fig 1A). However, the acceleration of missense divergence occurred in the North American population before the emergence of hyper-mutated viral lineages therein (Fig 1B, 4^th^ panel).

The hyper-mutated viral variants, which first emerged in immune-compromised patients with COVID-19 (Choi *et al*. 2020; Kemp *et al*. 2021), resembled the hyper-mutable microbial lineages with defective DNA repair systems that commonly emerged during LTEE studies (Lenski 2017). In LTEE populations, the “mutator” phenotype was maintained because the consistently higher benefits of new adaptive mutations out-weighing the cost of deleterious mutations in a controlled environment (Lenski 2017). Similarly, in an immune-deficient host environment, mutations that would have been deleterious in a normal host (e.g., those leading to hyper immunogenicity) may become neutral or beneficial to viral reproduction (Choi *et al*. 2020; Kemp *et al*. 2021). Freed from host immune constraints, viral evolution essentially follows the adaptive or mixed evolution models in which adaptive lineages dominate the viral population (Fig 1C and 1D, 3^rd^ *panel*). Nevertheless, all missense mutations observed in the hyper-mutated viral genomes are unlikely to be adaptive because neutral or slightly deleterious mutations are driven to high frequencies through genetic hitchhiking in an asexual population (i.e., genetic draft) (Gillespie 2000; Kim and Stephan 2000; Lang *et al*. 2013).

We note that, beyond adaptive mutations, the acceleration of missense divergence as measured by the *D_n_/D_s_* ratio could be caused by demographic forces. In the present study, we simulated viral evolution with a constant population size, although neutral and slightly deleterious missense mutations are expected to accumulate in the rapidly expanding viral populations (Messer and Petrov 2013). In addition, our analysis combined viral samples within a continent as representing a single population, whereas numerous local outbreaks and subsequent migrations between subpopulations are expected to contribute to increased viral genome diversity including missense divergence (see next section).

### Expanding genome diversity: demographic and selective causes

SARS-CoV-2 genomic diversity, measured by monthly average genome differences (*π*), increased in the six continents during the first year of the COVID-19 pandemic (Fig 2A). Expanding genomic diversity is expected for a nascent viral population before it reaches mutation-drift balance even if the population remains at a constant size (Fig 2C). Clearly, the global viral populations are far from reaching an equilibrium level of genomic diversity as the virus has spread within and across continents, mirroring the failures in local and global outbreak control. Furthermore, the increasing genomic diversity may be a reflection of increasing admixture of viral subpopulations distributed across the continents.

Beyond demographic forces, the relaxation of selective constraints and adaptive mutations may also have contributed to the rising viral genomic diversity. Ratios of missense to synonymous polymorphisms (*π_n_/π_s_*) were generally higher in continental populations than expected under strong purifying selection (Fig 2B), suggesting the accumulation of neutral and slightly deleterious missense mutations in the expanding viral population. Adaptive hypermutated lineages contributed to the increase in *π_n_/π_s_* ratios in later months in most continental populations and caused the elevated *D_n_/D_s_* ratios described in the previous section, An additional possible cause of rising viral genomic diversity is the presence of negative frequencydependent selection by which rare immune-escape variants gain a selectively advantage (Haven *et al*. 2011; Papkou *et al*. 2019).

### Asexuality, recurrent mutations, and recombination

We estimated the genome-wide levels of LD on the basis of 93 most frequent missense SNVs segregating in 8215 genomes sampled from six continents (Supplemental Material Table S3). These mutations were present with a frequency of 20% or higher in at least one month in one continent. Complete LD (*D*’ close to 1) dominated the *D*’ values between pairs of SNVs and, furthermore, there was no evidence of LD decay over genomic distances between the SNVs (Supplemental Material Fig S5). LD decay over distance is expected if recombination among viral strains occurs with sufficient frequency. In microbial species, recombination occurring at a rate comparable to the rate of point mutation is sufficient to cause LD decay over genomic distances (Fraser *et al*. 2007; Ansari and Didelot 2014). Thus we conclude that SARS-CoV-2 populations during the first year of pandemic were largely asexual with little evidence of recombination. The asexual population structure of SARS-CoV-2 populations mirrors the low recombination rates during previous SARS and Middle East respiratory syndrome (MERS) coronavirus outbreaks (Chinese SARS Molecular Epidemiology Consortium 2004; de Wit *et al*. 2016).

An analysis of SARS-CoV-2 genomes from early isolates suggested active recombination during human transmission based on a high level of homoplasy – independent mutations occurring at the same sites that cause inconsistencies with the viral phylogeny (Yi 2020). A prominent example of phylogenetically inconsistent mutation is the nonsynonymous SNV TTT[Phe]/TTG[Leu] at the genomic position 11083 of the *Nsp6* locus (Yi 2020). By reconstructing genome phylogeny through haplotype networks, we observed a similarly high level of homoplasy caused by either recombination or by mutations that have occurred independently in multiple evolutionary lineages (Supplemental Material Fig S6). Recurrent mutations and sequencing errors may have contributed to the observed homoplasy (Turakhia *et al*. 2020). Recurring mutations are inevitable in SARS-CoV-2 with its relatively small genome size. The chance of mutation recurrence increases as the pandemic spreads and persists. Indeed, we were able to estimate the rate of mutation recurrence with the use of *CovSimulator*. In a simulated population evolving under neutral conditions, ~2.9% genomic sites (860 out of the total of 29903 sites) experienced two or more mutations after 500 generations. This number was significantly greater than expected from a random Poisson process (*p*=2.1e-270 by a *χ^2^* test of goodness of fit). For two mutations occurring at the same genomic site, strong mutation biases seen during SARS-CoV-2 genome evolution stipulate a high chance of parallel base substitutions. For example, a mutation at a cytosine (C) site will almost certainly (with a ~95% chance) result in a thymine (T) (Table 2; Supplemental Material Fig S3).

It should be cautioned that a clonal population structure in SARS-CoV-2 does not imply an absence of or an inability of homologous recombination. In fact, coronaviruses are known for their high potential for homologous recombination in natural reservoirs as well as in the laboratory conditions (Masters 2006; Denison *et al*. 2011; Cui *et al*. 2019). SARS-CoV-2 genomes showed a mixed ancestry containing parts of the genome from coronaviruses associated with the pangolin (*Manis javanica*) and other parts from related viruses associated with the bat (*Rhinolophus affinis*) (Andersen *et al*. 2020; Lam *et al*. 2020). Consequently, the high clonality of SARS-CoV-2 populations is likely to be due to the explosive population growth worldwide (“epidemic structure”) rather than to an inability of recombination (Smith *et al*. 1993). By reducing clonal interference among competing adaptive mutations as well as by removing deleterious mutations without decreasing the frequencies of beneficial alleles, recombination is a powerful mechanism accelerating adaptation across species including microorganisms (Smith *et al*. 1993; Barton and Charlesworth 1998). As such, it is important to be vigilant about the rising chance of recombination among SARS-CoV-2 variants as the COVID-19 pandemic becomes entrenched. Previously, we have quantified recombination rates in natural populations of Lyme disease bacterium based on genome comparisons and computer simulations (Qiu *et al*. 2004; Haven *et al*. 2011). Similarly, *CovSimulator* can be used to detect the presence of recombination and to estimate recombination rates in SARS-CoV-2 populations through a comparison of homoplasy levels in populations simulated with and without recombination.

### Adaptation despite background selection: a model of SARS-CoV genome evolution

The highly clonal population structure of SARS-CoV-2 and the microbial species in LTEE studies implies that genetic variations across the entire genome are highly linked. As such, various selective forces interfere with one another including, for example, purifying selection at housekeeping loci, diversifying selection at antigenic loci, and adaptive evolution at host-binding sites (Hill and Robertson 1966; Gillespie 2000; Lang *et al*. 2013; da Silva and Galbraith 2017; Lenski 2017; Campos and Charlesworth 2019). In addition, neutral or even deleterious mutations may “ride along” with a newly emerged adaptive mutation and to reach high frequency in an asexual population.

On the basis of the conclusions on adaptive mutations contributing to genome variability and on the strong LD across the SARS-CoV-2 genome, we propose the mixed evolution as a model to understand the dynamics of SARS-CoV-2 genome evolution. In the mixed model (Figs 1C, 1D, 2C, and 2D), the majority of missense mutations were slightly deleterious (~80% probability with a multiplicative fitness cost of 0.95) and a small proportion of missense mutations were slightly adaptive (~10% probability with a multiplicative fitness benefit of 1.05).

First, we traced the genealogy of the final 20 sampled genomes, which showed a substantially shortened coalescence time since the most recent common ancestor (Fig 3A). Next, we tracked the frequencies of the top most frequent (≥5%) missense mutations for 500 generations (Fig 3B). Adaptive mutations (11 of a total of 19 mutations) dominated the final population. Nevertheless, not all fixed mutations were adaptive. Three neutral (#1, #2, and #4, in gray) and three deleterious missense (#2, #3, and #4 in blue) mutations became fixed through linkage with adaptive driver mutations, exemplifying genetic draft (Gillespie 2000). Conversely, not all adaptive mutations were destined to be fixed, indeed, one adaptive mutation (#5, in red) was lost because of competition with other adaptive mutations, exemplifying clonal interference (Lang *et al*. 2013; Maddamsetti *et al*. 2015). Thirdly, we generated the Muller diagrams, which grouped mutations sharing similar frequency trajectories (i.e., temporal linkage) into a single “genotype” (Fig 3C). The diagrams highlighted regular selective sweeps driven by adaptive mutations. Critically, it is clear from the Muller diagrams that within each “genotype” (e.g., G1, G2, G3, G8, and G9), at least one genetic change was the driver adaptive mutation. To facilitate comparison of evolutionary dynamics among evolutionary models, we provided the genome genealogies of the last-generation samples and the Muller diagrams of the topmost frequent missense mutations from all four models of simulated evolution as Supplemental Material Fig S4.

In summary, the mixed evolution model illustrates that, first, adaptive mutations and viral lineages quickly dominate the viral population despite that most of the missense mutations are deleterious. Second, neutral and deleterious mutations can become fixed through genetic hitchhiking with adaptive mutations. Third, adaptive mutations can be lost because of strong clonal interference. Fourth, recurring mutations become increasingly common because of a small viral genome, strong mutation biases, longer evolution time, and prolonged maintenance of adaptive lineages in the viral population. Fifth, temporal linkage among missense mutations provides a way to identify adaptive driver mutations. These conclusions are anticipated by observations from microbial LTEE studies as well as by results of theoretical analysis, both of which showed dominance of adaptive mutations in asexually evolving populations despite presence of strong purifying selection (Kim and Stephan 2000; Desai and Fisher 2007; Lenski 2017). We note that only one set of evolutionary parameters was used in the present simulation of the mixed model, the outcome of which would vary quantitatively with selection parameters including the proportions and strengths of deleterious and adaptive mutations.

### Spatiotemporal characteristics of adaptive mutations

The mixed model of SARS-CoV-2 genome evolution revealed a number of characteristics of adaptive mutations that are informative for their identification. First, adaptive mutations were over-represented in high-frequency SNVs (Fig 3). In one population simulated with mixed model, 258 adaptive mutations (10.9% out of a total of 2376 missense mutations) were present in the combined sampled genomes. However, 14 (45%) of the adaptive mutations were among the 31 missense mutations that have reached a frequency of 0.5% or higher. Second, the proportion of adaptive mutations among missense mutations increased over time (Fig 3). In the same simulated population, among the 20 genomes sampled from the last generation, the fixed missense mutations included 7 (70%) adaptive, 2 (20%) deleterious, and 1 (10%) neutral mutations. Third, in clusters of mutations that shared similar temporal trajectories, at least one of the consortium was the adaptive driver (e.g., G1, G2, G3, G8, and G9 in Fig 3). These characteristics of adaptive mutations suggest ways to identify adaptive mutations driving SARS-CoV-2 adaptation to humans through spatiotemporal tracking of mutation frequencies.

Guided by the above insights from the simulated evolution and LTEEs, we identified a genome-wide set of 101 missense mutations with a presence of 20% or higher frequency in at least one month within a continent (Supplemental Material Table S4). The high-frequency mutations were most often found in genes encoding the spike (S, *n*=21, 20.8%), nucleocapsid (N, *n*=19, 18.8%), Nsp3 (*n*=13, 12.9%), and ORF8 (*n*=9, 8.9%) proteins. Similarly, we identified a set of 52 missense mutations on the spike protein with a presence of 5% or higher frequency in at least one month within a continent (Supplemental Material Table S5). Mutations on the spike protein are of particular interest because of its use as vaccinogen. Half (*n*=26) of these spike protein mutations were located within the *N*-terminus domain (NTD) and receptor-binding domain (RBD), suggesting an oversized role the NTD and RBD mutations play in driving SARS-CoV-2 adaptation to humans. Sequences in NTD and RBD evolve faster relative to the genome average during coronavirus divergence, further supporting the role of mutations within these domains in driving viral adaptation to humans (Luk *et al*. 2019; Phan 2020; Cagliani *et al*. 2020) (Supplemental Material Fig S6).

We subsequently clustered these high-frequency mutations on the basis of their temporal trajectories within each continent. A heatmap of frequency trajectories of 52 missense mutations (with >5% frequency in at least one month) on the spike protein revealed clusters of mutations that were distributed either across the globe or more limitedly within continents (Fig 4). The globally distributed mutations included the D614G substitution that arose during the first month (January 2020) of the SARS-CoV-2 outbreaks in Asia and quickly reached global fixation. Clinical and experimental studies suggested enhanced human transmissibility but not increased disease severity associated with D614G viral variants (Korber *et al*. 2020; Volz *et al*. 2021; Plante *et al*. 2021). It is possible that missense mutations strongly linked with the D614G mutation, including P323L in Nsp12 (RNA polymerase) and R203K and G204R in the N (nucleocapsid) protein, may have also played a role in increased viral fitness in humans (Yang *et al*. 2020).

**Fig 4.**
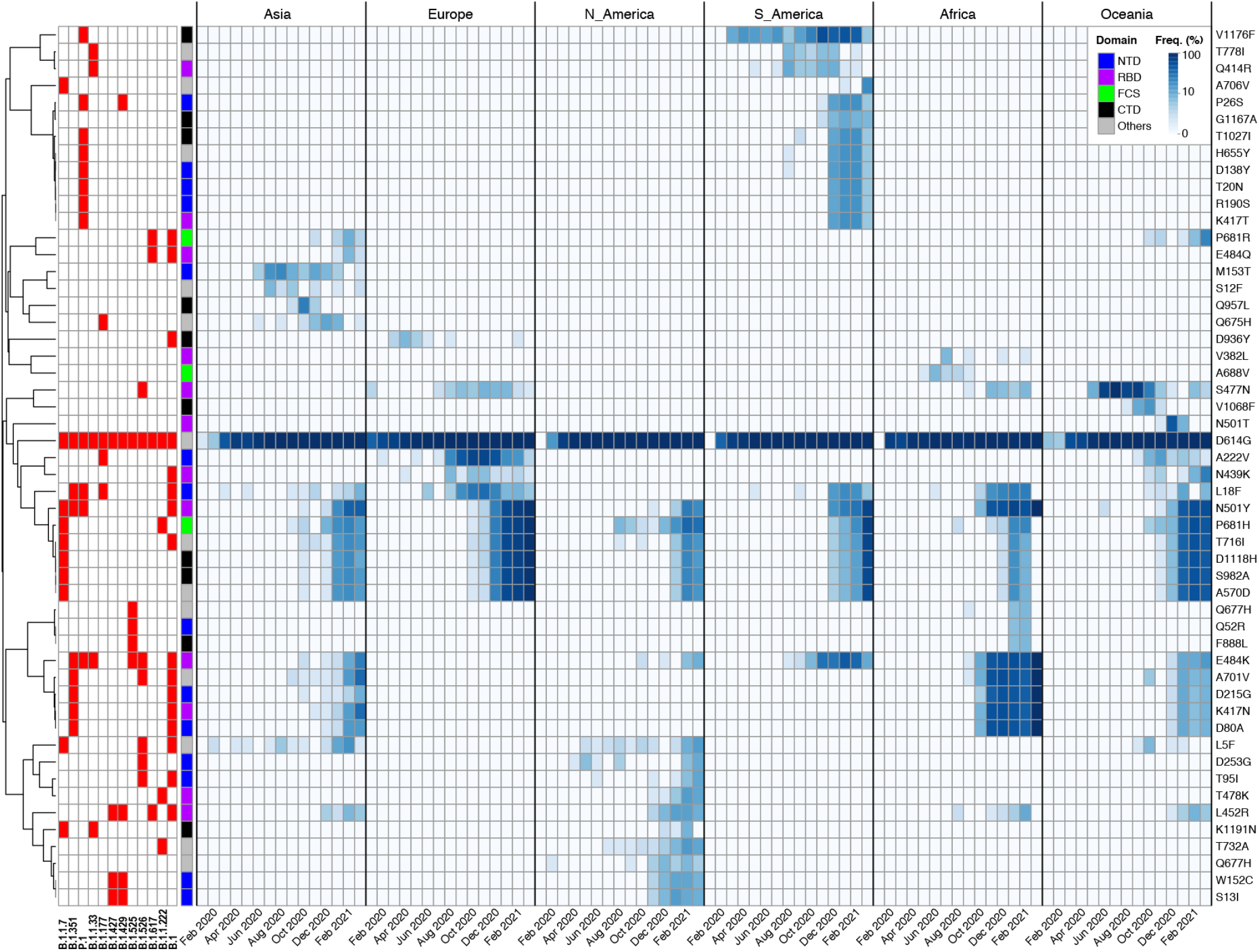
Frequency trajectories of high-frequency missense SNVs in six continental SARS-CoV-2 populations. The heatmap depicts allele frequencies (*colored cells*, in percentage, scaled with 10-based logarithm) by month (*columns*) of 51 missense mutations (*rows*) on the spike protein in viral populations from six continents (*vertical blocks*). Each mutation frequency was calculated on the basis of ~100 genomes randomly sampled from a month within a continent. Each mutation was present with a ≥5% allele frequency in at least one monthly sample. Mutations were grouped according to similarities in frequency trajectories (*rowside dendrogram*). The rowside table shows mutations associated with major viral lineages (Rambaut *et al*. 2020) (columns #1 through #12). Column #13 of the rowside table shows the spike protein domains associated with the mutations, including the N-terminus domain (NTD), receptor-binding domain (RBD), Furin cleavage site (FCS); and the C-terminus domain (CTD). The heatmap reveals the early rise and rapid fixation of the D614G mutation across the globe (dark blue stripe in the middle). Also discernable is rapid global spread of six spike protein mutations (N501Y, P681H, T716I, D1118H, S982A, and A570D) associated with the hyper-mutated B.1.1.7 lineage (rowside column #1) after its first emergence during October 2020 in Europe (Choi *et al*. 2020). Other mutations were associated with lineages that have so far shown limited geographic ranges, including the B.1.351 (originated in Africa, rowside column #2), P.1 (South America, rowside column #3), B.1.427 and B.1.429 (North American, rowside columns #6 and #7), and B.1.617 (Asia, rowside column #10) lineages. For early detection of human-adaptive mutations, it is necessary to track mutation frequencies at country and regional levels before they become more widespread (Fig 5).

A temporally linked group of six spike protein mutations – N501Y, P681H, T716I, D1118H, S982A, and A570D – associated with the hyper-mutated B.1.1.7 lineage (Fig 4) that emerged in September 2020 in England and quickly spread worldwide represent another set of mutations that have enhanced viral transmission in humans (Galloway 2021; Kemp *et al*. 2021). These spike protein mutations pose the additional risk of viral escape from protective immunity elicited with vaccines designed on the basis of the bat-adapted progenitor genome (Wang *et al*. 2021; Collier *et al*. 2021). Other mutations have so far been confined within one or more continents and have not reached global presence. These spike protein mutations included those associated with the B.1.351 lineage in Africa, the P.1 lineage in South America, the B.1.427/B.1.429 lineages in North America, and the B.1.617 lineage in Asia (Fig 4).

The strong candidates of human-adaptive mutations shown in the above have risen relatively early during the pandemic. The latest emergent human-adaptive mutations, however, would first reach high frequencies only in local outbreak populations. Thus, it is necessary to track allele frequencies at regional levels for early detection of human-adaptive mutations. As an example, we tracked the spatiotemporal frequencies of 56 spike missense mutations with ≥5% allele frequencies in at least one month in the United States and and its five states including Washington, California, New York, Texas and Michigan (Fig 5). Except for the globally fixed D614G mutations and mutations associated with the B.1.1.7, B.1.427 and B.1.429 lineages, mutations associated with the latest emergent viral lineages only reached the threshold 5% level within the states and not at the national level. For example, the B.1.526 and B.1.243 lineages emerged during December 2020 in New York and have not yet spread to the other four states. The B.1.2 lineage in Washington and the B.1.234 lineage in Michigan have thus far been observed only within the states.

**Fig 5.**
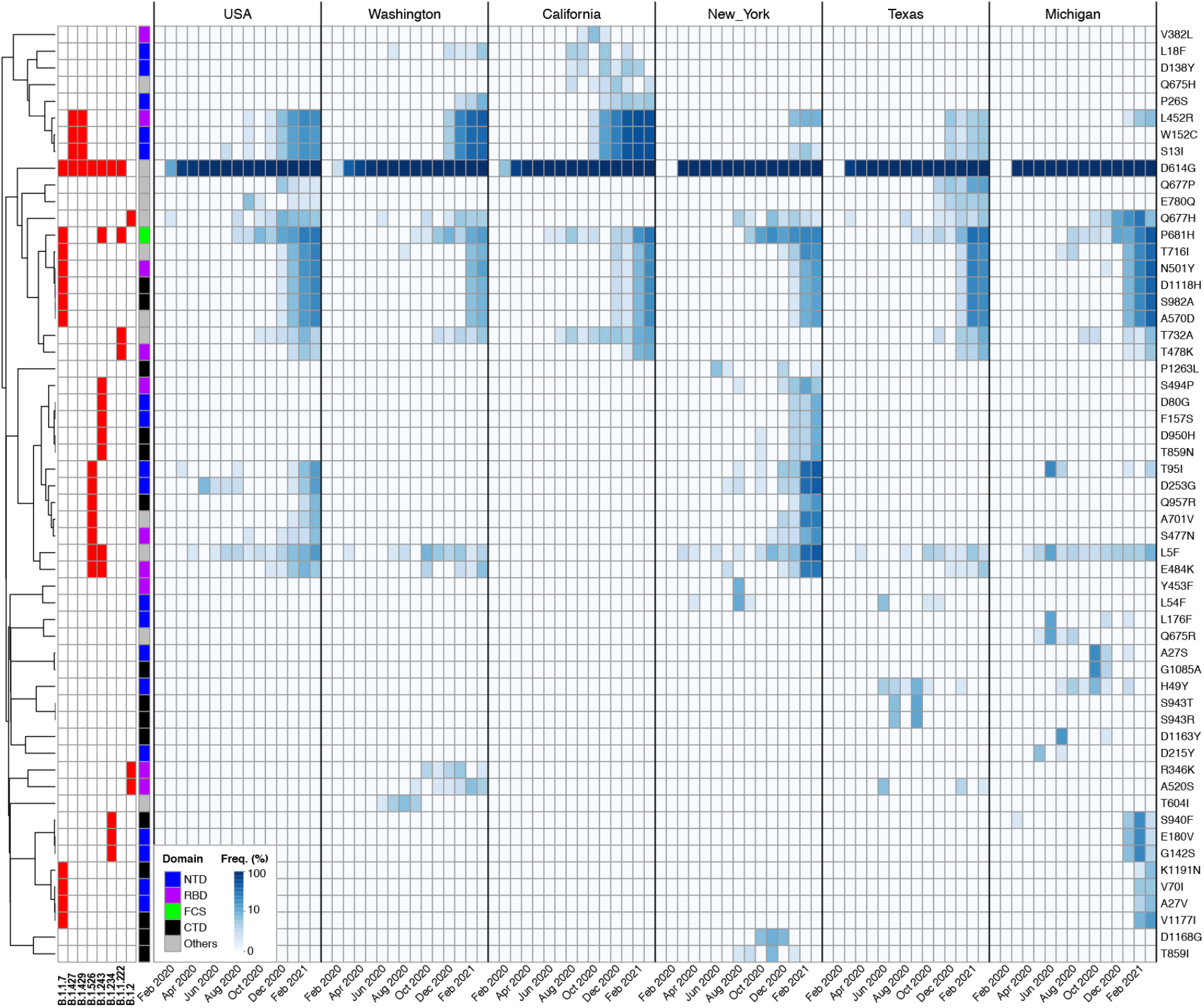
Tracking emergent adaptive mutations at regional levels. The heatmap depicts monthly (*columns*) frequencies (*colored cells*) of 56 missense mutations (*rows*) on the spike protein that were present with ≥5% allele frequencies in at least one monthly SARS-CoV-2 sample from the United States and its five states (*vertical blocks*). As in the global heatmap (Fig 4), the D614G mutation reached fixation across the country since March 2020. The European lineage B.1.1.7 (*rowside column* #1) first arrived the US in December 2020 and quickly spread to all five states. The B.1.427 and B.1.429 lineages (*rowside columns* #2 and #3) were first identified in fall 2020 in California and have spread to the four other states by the end of March 2021. Similarly, the B.1.1.222 (*rowside column* #7) lineage was first identified in summer in California and has since spread to Washington, Texas, and Michigan. The B.1.526 and B.1.243 (*rowside columns* #4 and #5) lineages emerged during December 2020 in New York and have not yet spread to the other four states. Similarly, two other lineages have thus far not yet spread outside of the state of origination, including the B.1.2 lineage (rowside column #8) in Washington and the B.1.234 lineage (rowside columns #6) in Michigan. None of the latter four regional lineages has reached a ≥5% frequency at the national level (1^st^ vertical block), highlighting the importance of identifying human-adaptive mutations by tracking mutation frequencies at the level of local populations.

In summary, whereas missense mutations in the SARS-CoV-2 genome that have reached high frequency at local or global levels are not necessarily human-adaptive mutations because of the possibilities of genetic drift and hitchhiking, clusters of missense mutations that display temporal linkage and have reached high frequencies are indicative of adaption to humans. Within each of the cluster of temporally linked high-frequency missense mutations, we expect at least one to be a human-adaptive driver mutation. As such, the high-frequency clusters of missense mutations are top-priority candidates for clinical development of therapeutics and vaccines that target human-adapted viral variants.

### Concluding remarks

In the present work, we used realistic simulations of genome evolution and insights from microbial long-term evolution experiments (LTEEs) (Tenaillon *et al*. 2016; Cvijović *et al*. 2018) to understand the evolutionary transition of the SARS-CoV-2 virus from a bat-adapted to a human-adapted pathogen. The two evolving systems share salient evolutionary characteristics including strong purifying selection associated with a compact genome and large population sizes, forced adaptation to a new environment, and an asexual population structure. Not surprisingly, the variety of adaptive dynamics occurred in LTEEs were all discernable during SARS-CoV-2 evolution including the early rise and rapid fixation of adaptive mutations, clonal interference with competing adaptive mutations, fixation of neutral and deleterious mutations due to genetic hitchhiking. Specifically, both LTEEs and our analysis suggest that temporal linkage among mutations is a sensitive means for identifying emerging human-adaptive mutations and vaccine-escape mutations, particularly when mutation frequencies are tracked at the local and regional levels.

Epidemiological models based on human coronaviruses and influenza viruses predict the COVID-19 to be a recurrent seasonal disease in the next 2 ~ 5 years (Kissler *et al*. 2020; Cobey 2020). We expect to see continued expansion of viral genome diversity as the pandemic persists, entailing increasing risks for viral adaptation to humans and viral escape from natural and vaccine-induced protective immunity. Prolonged pandemic incurs the additional risks of recurrent mutations and recombination among viral variants, which would accelerate viral adaptation to humans. The software systems we developed facilitate real-time tracking of SARS-CoV-2 outbreaks. The *CovSimulator* software system is capable of modeling the trajectories of SARS-CoV-2 genome evolution and could be further improved by including more realistic parameters such as population expansion, migration and admixture between subpopulations, and frequency-dependent fitness imitating vaccine-escape mutations. The second software system associated with the *cov-db* genome database is capable of rapid tracking of emergent adaptive mutations through temporal sampling of genomes in a continent, country, or region therein. Thirdly, the cov.genometracker.org website provides a public-friendly user interface to search, browse, and visualize SARS-CoV-2 genome evolution and mutation trajectories (Supplemental Material Fig S7). Mutations appear in genes encoding proteins that down-regulate host immune responses (e.g., ORF3a and ORF8) and bind host cells (e.g., Spike) are high priority targets for the development of therapeutics and vaccines against human-adapted SARS-CoV-2 variants.

## List of Supplemental Material

- Supplemental Tables

○ Table S1. Estimated total number of synonymous and nonsynonymous sites
○ Table S2. McDonald and Kreitman analysis
○ Table S3. LD statistics between pairs of 91 high-frequency SNVs
○ Table S4. High-frequency SNVs with per-site rates and homoplasy indices
○ Table S5. High-frequency SNVs on the spike protein
- Supplemental Figures

○ Fig S1. The CovSimulator software system
○ Fig S2. The cov-db software system
○ Fig S3. Genome base composition and substitution rates
○ Fig S4. Genome genealogies and mutation trajectories in simulated evolution
○ Fig S5. Genome-wide linkage disequilibrium
○ Fig S6. Per-site rates, homoplasy, and haplotype network of spike protein mutations
○ Fig S7. Web-interactive mutation tracker (on cov.genometracker.org)

## Acknowledgements, Funding and Conflicts of Interest

We gratefully acknowledge the authors and the laboratories that originated and submitted the sequences to the GISAID’s EpiCoV™ Database on which this research was based. We thank Christopher Panlasigui for participating in discussions and Dr Yozen Hernandez for computer system administration. S.A. and L.L. are supported in part by the Graduate Program in Biology from the Graduate Center, City University of New York.

This work was supported in part by the National Institute of Allergy and Infectious Diseases (NIAID, AI139782, to W.Q.) and National Institute of Biomedical Imaging and Bioengineering (NIBIB, EB030275, to Brian Zeglis and W.Q.) of the National Institutes of Health (NIH) of the United States of America.

The authors declare no conflicts of interest.

## Author Contributions

S. Akther initiated the project, performed all evolutionary analysis, and participated in manuscript drafting. E. Bezrucenkovas downloaded viral genomes, prepared associated metadata, and developed the prototype evolution simulator. L. Li participated in the development of the simulator and performed simulation, linkage, and evolutionary rate analyses. B. Sulkow contributed to conceptural development and performed Muller diagram analysis. L. Di developed the genome database and web browser. D. Pante contributed to haplotype network and mutation analyses, C.L. Martin contributed to mutation analysis. B.J. Luft contributed to conceptual development and manuscript preparation. W. Qiu designed the software systems, implemented the simulator, and drafted the manuscript.

